# Directed evolution of *Escherichia coli* with lower-than-natural plasmid mutation rates

**DOI:** 10.1101/301382

**Authors:** Daniel E. Deatherage, Dacia Leon, Álvaro E. Rodriguez, Salma K. Omar, Jeffrey E. Barrick

## Abstract

Unwanted evolution of designed DNA sequences limits metabolic and genome engineering efforts. Engineered functions that are burdensome to host cells and slow their replication are rapidly inactivated by mutations, and unplanned mutations with unpredictable effects often accumulate alongside designed changes in large-scale genome editing projects. We developed a directed evolution strategy, <u>P</u>eriodic <u>Res</u>election for <u>E</u>volutionarily <u>R</u>eliable <u>V</u>ariants (PResERV), to discover mutations that prolong the function of a burdensome DNA sequence in an engineered organism. Here, we used PResERV to isolate *E. coli* cells that replicate ColE1 plasmids with higher fidelity. We found mutations in DNA polymerases I and IV and in RNase E that reduce plasmid mutation rates by 6-to 30-fold. The PResERV method implicitly selects to maintain the growth rate of host cells, and high plasmid copy numbers and gene expression levels are maintained in some of the evolved *E. coli* strains, indicating that it is possible to improve the genetic stability of cellular chassis without encountering trade-offs in other desirable performance characteristics. Utilizing these new antimutator *E. coli* and applying PResERV to other organisms in the future promises to prevent evolutionary failures and unpredictability to provide a more stable genetic foundation for synthetic biology.

## INTRODUCTION

Populations of cells engineered to function as factories or biosensors experience a failure mode that is peculiar to living systems: they evolve. Unwanted evolution is a foundational problem for bioengineering that limits the efficiency and predictability of metabolic and genome engineering efforts (1–5). Often an engineered function diverts critical resources from cellular replication or otherwise interferes with growth or homeostasis (6, 7). In these cases, ‘broken’ cells with mutations that inactivate the engineered function can rapidly outcompete the original design (8–10). The rate at which an engineered function decays within a cell population in this manner can be summarized as an evolutionary lifetime or half-life (8) or defined in terms of an evolutionary landscape by the rates at which various mutational failure modes occur and their respective fitness benefits (9, 10).

It is sometimes possible to edit a genome to eliminate or reduce the rate at which certain types of mutations occur (11–14) or to devise a way of reducing the burden of an engineered function (2, 15). However, given the complexity of DNA replication and repair processes and the multifarious ways that an engineered function can burden a host cell, a point is generally reached at which it is difficult to further improve upon the reliability of a cell. Directed evolution is an effective strategy for optimizing the performance of complex systems with many interacting components, even when they include unknown factors. For example, it has been used to engineer novel enzymes that outperform their natural counterparts (16) and to tune artificial gene circuits to effectively perform logic operations (17).

Given the similarly complex constraints underlying cellular mutagenesis and the fitness burdens of diverse engineered functions, we reasoned that a directed evolution procedure, <u>P</u>eriodic <u>Res</u>election for <u>E</u>volutionarily <u>R</u>eliable <u>V</u>ariants (PResERV) (**Fig. 1**), could be an effective strategy for improving the evolutionary reliability of an engineered cell. In PResERV, one artificially selects for mutant cells that exhibit improved maintenance of a burdensome engineered function over tens to hundreds of cell divisions. We expected that PResERV might isolate cells with mutations that either reduced the rate at which failure mutations occurred or the fitness burden of the engineered function, or both, possibly in ways that would generalize to stabilizing other engineered functions in the evolved cells. Here, we describe *E. coli* strains evolved by PResERV that exhibit lower-than-natural mutation rates for genes encoded on high-copy plasmids, thereby stabilizing them against unwanted evolution.

**Figure 1.**
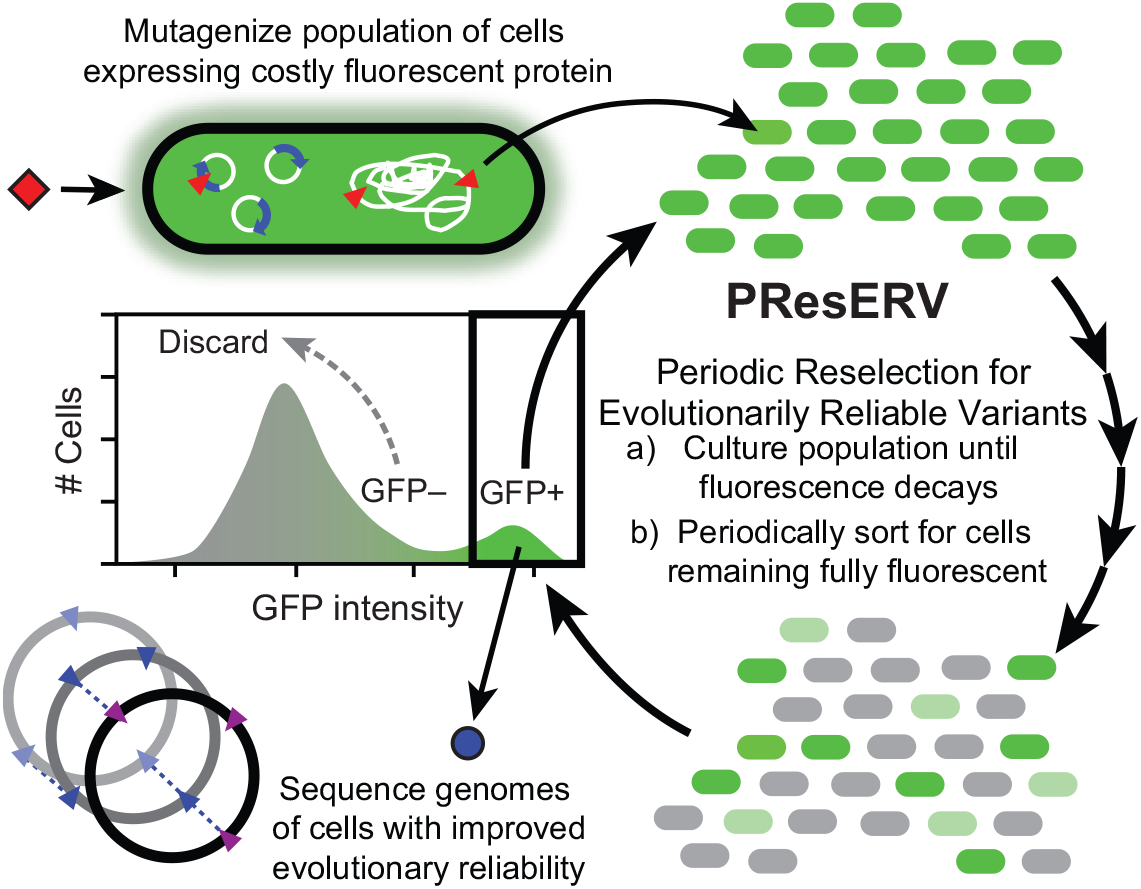
Periodic Reselection for Evolutionarily Reliable Variants (PResERV) method. PResERV begins with a cell expressing GFP to such a high level that it imposes a significant fitness burden. After mutagenesis, the cell population is cultured through enough cell doublings that mutants with reduced GFP expression take over the majority of the population. Periodically, the population is sorted to retain only those cells that remain as fluorescent as the original strain, enriching for mutant host cells with reduced mutation rates or a lower fitness cost for GFP expression. Once the evolutionary stability of GFP expression increases, fluorescent cells are isolated and their genomes are sequenced to identify and characterize the genetic changes that contribute to this improvement. Regrowth of cells during PResERV implicitly selects for only those mutants that achieve improved genetic stability without introducing any trade-offs that significantly reduce cellular growth rates.

## MATERIAL AND METHODS

### Culture conditions

*E. coli* was grown as 10 mL cultures in 50 mL Erlenmeyer flasks with incubation at 37°C and 120 r.p.m. orbital shaking over a diameter of 1 inch unless otherwise noted. The Miller formulation of Lysogeny Broth (LB) was used (10 g/L tryptone, 5 g/L yeast extract, and 10 g/L NaCl). Media were supplemented with 100 μg/mL carbenicillin (Crb), 20 μg/mL chloramphenicol (Cam), 50 μg/mL kanamycin (Kan), 100 μg/mL rifampicin (Rif), and 1 mM Isopropyl β-D-1-thiogalactopyranoside (IPTG), as indicated. Bacterial cultures were frozen at −80°C after adding glycerol as a cryoprotectant to a final concentration of 13.3% (v/v).

### Strains and plasmids

The progenitor strain (BW25113) of the Keio knockout collection (18) was transformed with pSKO4. This plasmid encodes the redesigned I7101 (R0010+E0240) circuit that was edited to remove unstable repeat sequences in a prior study by Sleight et al. on the BioBrick cloning vector pSB1A2 backbone (19). It is a ColE1 group plasmid with a pBR322 origin of replication. Plasmid pTEM-1.D254tag encodes β-lactamase with the codon for an amino acid at a surface-exposed position in its structure at which multiple amino acid substitutions are compatible with enzyme function replaced with a TAG stop codon (20). It has a pBR322 origin of replication and also encodes the *rop* protein.

### UV mutagenesis

BW25113 cells containing pSKO4 were cultured overnight to stationary phase in LB-Crb. Then, these cultures were pelleted by centrifugation and resuspended in an equal volume of sterile saline. Eleven 120 μl droplets of these cell suspensions were spotted on petri dishes and subjected to 27,500 μJ/cm^2^ of 254 nm UV radiation in a UVP CL-1000 crosslinker. After UV exposure, 100 μl from each droplet was combined and pelleted by centrifugation to collect ~2.5×10^6^ surviving cells. These cells were inoculated into 10 mL of LB-Crb and grown to a final density of ~2×10^9^ cells/ml. This mutagenized library was archived as a frozen stock.

### PResERV directed evolution procedure

All growth steps were conducted in 10 mL of LB-IPTG-Crb. We used 0.1 mL of the mutagenized library to found the experimental population. After overnight growth to saturation, we propagated the population through daily 1:1000 dilutions of saturated cultures into fresh media followed by regrowth. GFP expression was monitored using a BD Fortessa flow cytometer. Periodically, overnight cultures were diluted to ~2.5×10^6^ cells/mL in HPLC grade water and stained with 150 nM of the nucleic acid dye SYTO 17 (Life Technologies). The GFP+ portion of the population was calculated as the percentage of SYTO 17 positive cells with at least the ancestral level of GFP fluorescence by flow cytometry. When only ~15-25% or fewer of cells in the population were GFP+, instead of a normal transfer, the population was diluted to ~2.5×10^6^ cells/mL in HPLC grade water and between ~4×10^4^ and ~5×10^5^ GFP+ cells were sorted into 10 mL of fresh LB-Crb using a BD FACSAria IIIu. The SYTO 17 dye was found to decrease cell viability, so we did not use gating on this counterstain in these sorting steps, at the cost of less efficient enrichment of GFP+ over GFP− cells.

### Isolation of evolved cells and plasmids

The evolved population was plated on LB-IPTG-Crb and six visibly GFP+ colonies were selected at random for further study. Each of these clonal isolates was grown overnight in LB-IPTG-Crb before isolating its plasmid and creating a frozen stock. To select for plasmid loss, we also diluted these cultures 1:1000 into media lacking Crb (LB-IPTG). After overnight growth, dilutions of the resulting cultures were then plated on LB-IPTG agar. GFP– colonies were patched onto LB-Crb agar to ensure the lack of fluorescence was caused by loss of plasmid rather than a GFP mutation. One colony which had been cured of its plasmid was selected from each of the original six evolved clones and retransformed with the wild-type plasmid. The wild-type BW25113 strain was separately transformed with the plasmids isolated from each of the six evolved clones. GFP decay experiments were carried out on the resulting twelve strains to examine them for evidence of increased evolutionary stability.

#### GFP decay curves

For each of the twelve strains tested, individual colonies were used to inoculate nine replicate *E. coli* populations. In order to more accurately estimate the number of cell doublings elapsed since the single-cell bottleneck, care was taken to transfer all cells in each colony to the first liquid culture by excising and transferring the piece of agar underneath and around each colony. Each population was then subjected to daily transfers under the same conditions as the PResERV experiment, while monitoring GFP fluorescence using SYTO17 staining and the BD Fortessa flow cytometer as describe above. For creating graphs of the percentage of cells remaining GFP+ over time, flow cytometry data were analyzed in R using the flowCore Bioconductor package (v1.42.3) (21). Among the events successfully exhibiting a SYTO17 signal, cells were classified as GFP+ if they were above a uniform signal intensity across all samples in an experiment that was determined by examining the wild-type distribution. For graphing and comparing the initial GFP fluorescence in each strain, median intensity values for the GFP+ subpopulation were log2 transformed before performing statistical analyses.

### Genome sequencing

DNA was extracted from stationary phase *E. coli* cultures using the PureLink Genomic DNA Mini kit from Life Technologies. Purified DNA was fragmented using the Covaris AFA system and Illumina sequencing libraries were prepared using the NEBNext DNA Library Prep Reagent Set for Illumina kit from New England Biolabs. An Illumina HiSeq 2500 was used to generate 2 ×125 paired-end reads from each sample at The University of Texas at Austin Genome Sequencing Analysis Facility (GSAF). FASTQ files have been deposited in the NCBI sequence read archive (SRP090775). Mutations in each of the evolved strains were predicted by using *breseq* (version 0.32.0a) (22) to compare the Illumina reads to the *E. coli* BW25113 reference genome (GenBank:CP009273.1). Genetic differences between this reference sequence and all four sequenced samples were assumed to have existed in the ancestral *E. coli* strain used to initiate the PResERV experiment and are not reported.

### Strain reconstruction

Donor strains from the i-Deconvoluter library (23) were used to revert the three candidate evolved alleles we tested in two evolved clones (AER8 and AER12) back to wild type sequences using P1 transduction as previously described (24), except that only 2 μl of lysate was used in each transduction. Lysate from SMR20954, SMR20794, and SMR20838 was combined with the appropriate evolved strain with a mutation in *polA, polB*, or *rne*, respectively, and plated on LB agar plates containing Kan. Resultant colonies were screened for correct replacement via Sanger sequencing before FLP recombinase was used to remove the linked Kan^R^ cassette used for selection of transductants as previously described (25). The strain with *rne* reverted to the wild-type allele was subjected to a second round of P1 transduction to also revert the *polB* mutation, and the Kan^R^ cassette was again removed via FLP recombination.

### Mutation rate measurements

Luria-Delbrück fluctuation tests were carried out to measure mutation rates (26). For plasmid mutation rates, strains cured of plasmid pSKO4 were transformed with plasmid pTEM-1.D254tag after any genetic modifications to modify evolved alleles. Cultures were grown in LB-Cam to select for retention of this plasmid and plated on LB-Cam agar additionally supplemented with 500 μg/ml Crb to select for mutants. Thus, this assay measures the aggregate rate of all mutations that revert this stop codon to a permitted sense codon. For chromosomal mutation rates, LB agar containing 100 μg/ml rifampicin (Rif) or 60 μg/ml D-cycloserine (DCS) was used for the selective conditions. Rif resistance requires specific point mutations in the *rpoB* gene (27). In minimal media DCS resistance requires a loss-of function mutation specifically in the *cycA* gene (28), but mutations in additional targets may also be possible in the rich LB media used here.

Crb^R^ plasmid and Rif^R^ chromosomal mutation rates were determined by taking an overnight culture of a strain and transferring ~1,000 cells from a dilution in sterile saline to multiple 0.2 mL test tubes containing non-selective media. After growth of the replicate test tube cultures overnight to saturation, the entire volume of each one was either plated on a selective LB agar plate or a dilution was plated on a non-selective LB agar plate. A total of 6 non-selective and 48 or 12 selective plates were used for plasmid and chromosomal mutations respectively. Mutant colonies on the selective plates were counted after 48 h of incubation. For DCS chromosomal mutation rates, 1.0 mL LB cultures and 12 selective media plates were used. Additionally, only a portion (25 μl) of these cultures were plated on the selection LB agar. Colony counts on selective and non-selective plates were used to estimate mutation rates using the rSalvador R package (v1.7) (29). We used its likelihood ratio methods for calculating confidence intervals and the statistical significance of differences between mutation rate estimates.

### Plasmid copy number determination

Cells containing the pTEM-1.D254tag plasmid were revived in LB-Cam with overnight growth from frozen stocks and then diluted 1000-fold into fresh media and harvested at exponential phase (OD600 ~0.5). Three cell pellets from different biological replicates were collected for each strain. Primer pairs were designed to amplify products from either the antibiotic resistance gene (*cat*) in the pTEM-1. D254tag plasmid or the *ftsZ* gene in the *E. coli* chromosome. The *cat* primers were 5’-GTGAGCTGGTGATATGGGATAG and 5’-CCGGAAATCGTCGTGGTATT. The *ftsZ* primers were 5’-GCAAGGTATCGCTGAACTGA and 5’-CGTAGCCCATCTCAGACATTAC. Two amplification reactions for each of the two primer pairs were conducted using Power SYBR Green PCR Master Mix reagents and a ViiA7 Real-Time PCR System (ThermoFisher). Plasmid copy number per chromosome was estimated using the ΔΔCt method after averaging technical replicates for each biological replicate (30). Graphing and statistical analyses were performed using log2 transformed values of copy number estimates for each biological replicate. The mean copy number estimated for the pTEM-1.D254tag plasmid in the wild-type BW25113 strain was 410.

## RESULTS AND DISCUSSION

### PResERV experiment with a ColE1 plasmid in *E. coli*

We applied PResERV to *E. coli* K-12 strain BW25113 (18) transformed with pSKO4, a high-copy-number pBR322 plasmid (ColE1-type origin) that encodes GFP under control of an inducible promoter (19). GFP expression is a generic proxy for a costly engineered function in this scenario. A UV-mutagenized library of cells containing pSKO4 was propagated through daily 1:1000 serial transfers in the presence of antibiotic selection for plasmid retention. Under these conditions, cells with mutations in pSKO4 that inactivate or reduce costly GFP expression evolve, outcompete fully fluorescent cells, and constitute a majority of the population within a few days (19). The fluorescence of cells in the PResERV population was periodically monitored by flow cytometry. When ~75% or more of the cells exhibited reduced GFP signal, cell sorting was used to isolate ~10^5^ cells that remained as fluorescent as the ancestor to continue the population. We subjected this population to a total of 8 sorts spread throughout 30 regrowth cycles (**Fig. 2A**).

**Figure 2.**
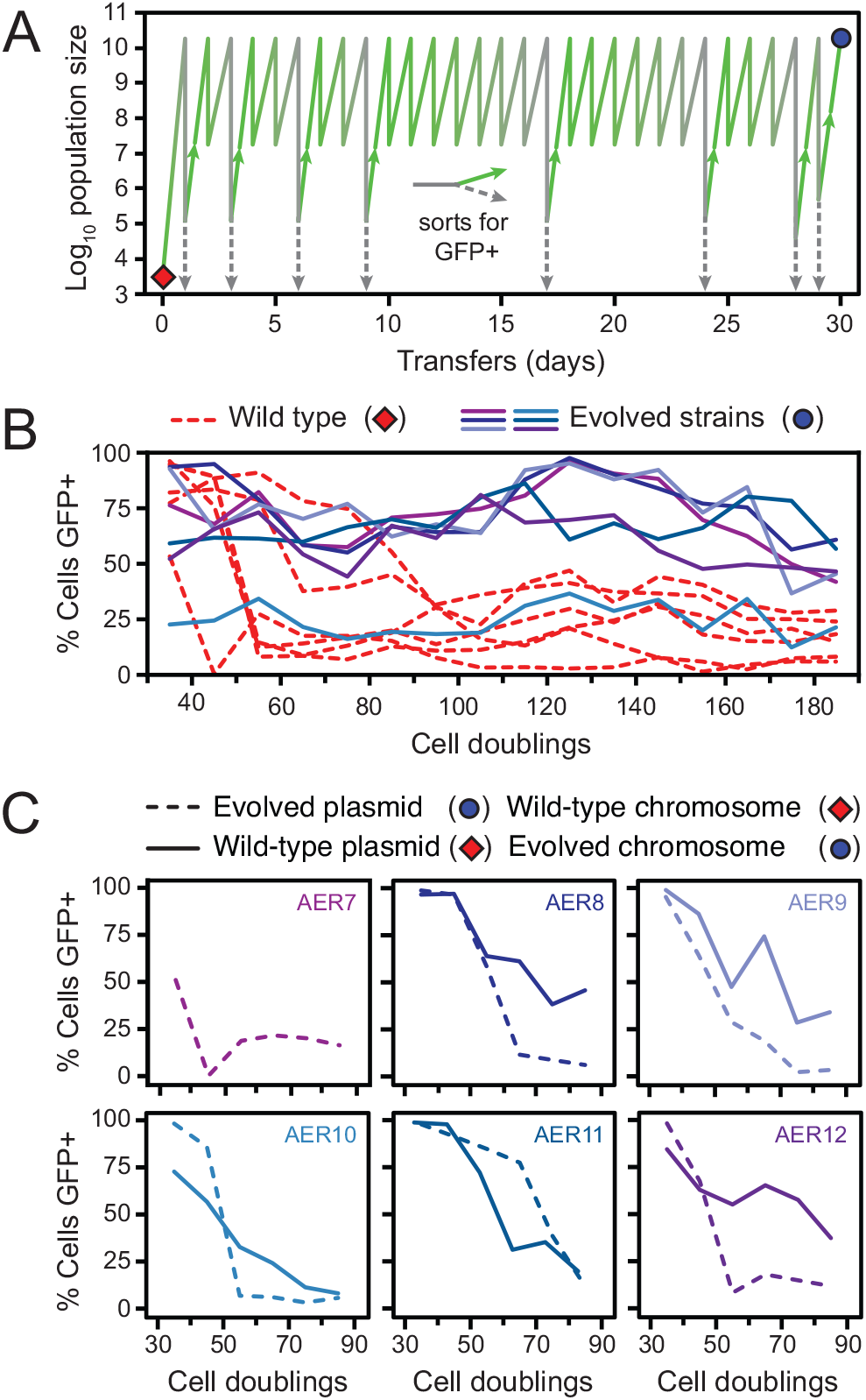
PResERV applied to an *E. coli* plasmid. (**A**) Propagation and sorting regimen used to perform PResERV on an *E. coli* population in which GFP was expressed from plasmid pSKO4, a high-copy plasmid with a pBR322 origin of replication. The red diamond denotes the wild-type strain that was UV-mutagenized prior to beginning PResERV. Dashed grey and solid green bifurcating lines show when the population was sorted to retain fully fluorescent cells (GFP+). The blue circle indicates when fluorescent clones were isolated and sequenced. (**B**) Populations initiated from six clones isolated from different populations at the end of PResERV (blue and purple solid lines) were allowed to evolve alongside six replicates of the non-mutagenized, wild-type *E. coli* strain-plasmid combination (red dashed lines). Cells were considered GFP+ if they maintained a fluorescent intensity above a threshold level that was kept constant across all tested strains. (**C**) For each evolved PResERV strain, its plasmid was isolated and transformed into the wild-type strain containing no plasmid (dashed lines), and the evolved strain was cured of its plasmid and re-transformed with the wild-type pSKO4 plasmid (solid lines). Populations initiated from these strains were propagated and monitored as in **B**. In most cases, the evolved chromosome rather than the plasmid seemed to be responsible for all or most of the improved reliability. The stability of AER7 re-transformed with the wild-type plasmid was not determined because of difficulty curing the plasmid from this strain. The same colors are used for each PResERV strain in panels B and C.

Six *E. coli* clones designated AER7–AER712 were isolated from the final population for further characterization. Five of these maintained more fully-fluorescent cells for more cell doublings than the unevolved wild-type strain with the wild-type pSKO4 plasmid (**Fig. 2B**). Mutations in the plasmid, the *E. coli* chromosome, or both could have been responsible for these improvements. To determine which was the case, we cured these cells of their plasmids and retransformed them with the wild-type pSKO4 plasmid, and we also isolated plasmids from each of the evolved strains and transformed them into unevolved wild-type *E. coli* cells. For four of these strains (AER7, AER8, AER9, and AER12), the improvement in the evolutionary lifetime of GFP expression appeared to be mainly associated with mutations located on the *E. coli* chromosome (**Fig. 2C**).

### Mutations in PResERV strains

We sequenced the genomes of these four evolved clones to understand the genetic basis of their improved reliability (**Fig. 3**). In agreement with the re-transformation tests, no mutations were found in the pSKO4 plasmid in any of these strains. Each contained from four to ten mutations in the *E. coli* chromosome. These mutations could theoretically lead to the improved maintenance of GFP expression that we observed by reducing the burden of GFP expression from the plasmid or by reducing the rate at which mutations that inactivate GFP arise. Therefore, we examined the lists of mutations in these strains to see if they hit any genes known to be involved in these processes.

**Figure 3.**
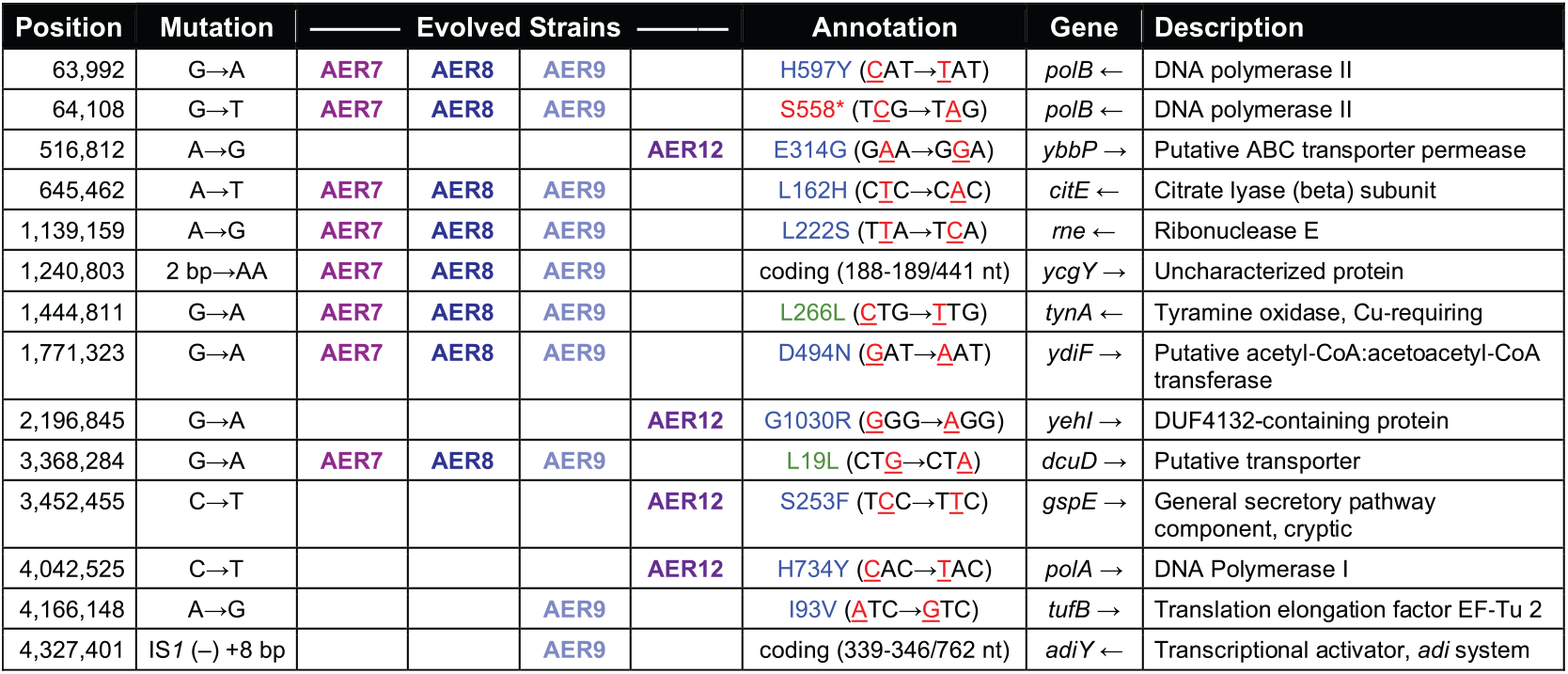
Mutations in PResERV strains. The genomes of four evolved strains were re-sequenced to identify mutations that accumulated during the directed evolution procedure. The position column shows the coordinate of the first affected base pair defined relative to the *E. coli* K-12 BW25113 genome (GenBank:CP009273.1). The mutation column shows base changes on the top strand of the genome, except for the IS*1* element in AER9 that inserted in the reverse direction and duplicated bases 4,327,401–4,327,408 at the target site on each side of the new IS copy. The annotation column shows the amino acid changes and codon changes caused by single-base substitutions or the locations of bases affected within a gene for other mutations. The gene column includes arrows showing the genomic strand on which each mutated gene is located.

Two of these strains (AER7 and AER8) had eight identical mutations while a third strain (AER9) had these same eight mutations plus two additional ones. All three shared mutations in *polB* and *rne* that were candidates for affecting evolutionary stability. PolB (Pol IV) is a stress-induced error-prone DNA polymerase that lacks proofreading activity. The *polB* gene sustained two mutations in these PResERV strains: a missense mutation (H597Y) and a nonsense mutation earlier in the reading frame (S558*). Deletion of *polB* in the clean-genome *E. coli* strain MDS42 has been shown to lead to ~30% lower chromosomal mutation rates (12). The *rne* gene (encoding RNase E) contains a missense mutation (L222S) in all three strains. RNase E regulates the copy number of ColE1 origin plasmids in *E. coli* by processing the RNAI antisense regulator of the RNAII replication primer (31). Mutants defective in *rne* accumulate higher levels of RNA I and have reduced plasmid copy number (32).

The fourth sequenced strain (AER12) had a completely different set of four mutations, which included a missense mutation in *polA* (H734Y). PolA (Pol I) is the DNA polymerase that is utilized primarily for filling gaps during lagging strand synthesis and in DNA repair in *E. coli*. Antimutator variants of PolA that lower the frequencies of mutations observed on a ColE1 reporter plasmid have been identified previously by screening a sequence library created by mutagenizing a PolA variant lacking 3’→5’ exonuclease proofreading activity (33). An H734Y substitution was also observed among the 592 polymerase variants tested in that study.

### Evolutionary stability and mutation rates in PResERV and reconstructed strains

To test whether these three mutations contributed to the increased evolutionary reliability of the PResERV strains, we tested *E. coli* strains in which we reverted the evolved alleles back to their wild-type sequences. We then propagated multiple populations of wild-type *E. coli*, two evolved clones (the clone with the *polA* mutation and one of the three clones containing mutations in *polB* and *rne*), and four revertant strains (one for each mutation and also a strain in which *polB* and *rne* were both reverted) under the same conditions as the initial evolution experiment and monitored the loss of fluorescence over the course of ~100 cell doublings (**Fig. 4A**). Mutations in *polA* and *rne* appeared to be responsible for most or all of the improved stability, as reverting these mutations reduced the evolutionary lifetime of GFP expression back to a level similar to that observed in the wild-type strain. In contrast, reverting the *polB* mutation alone or in combination with reverting *rne* did not appreciably affect GFP stability.

**Figure 4.**
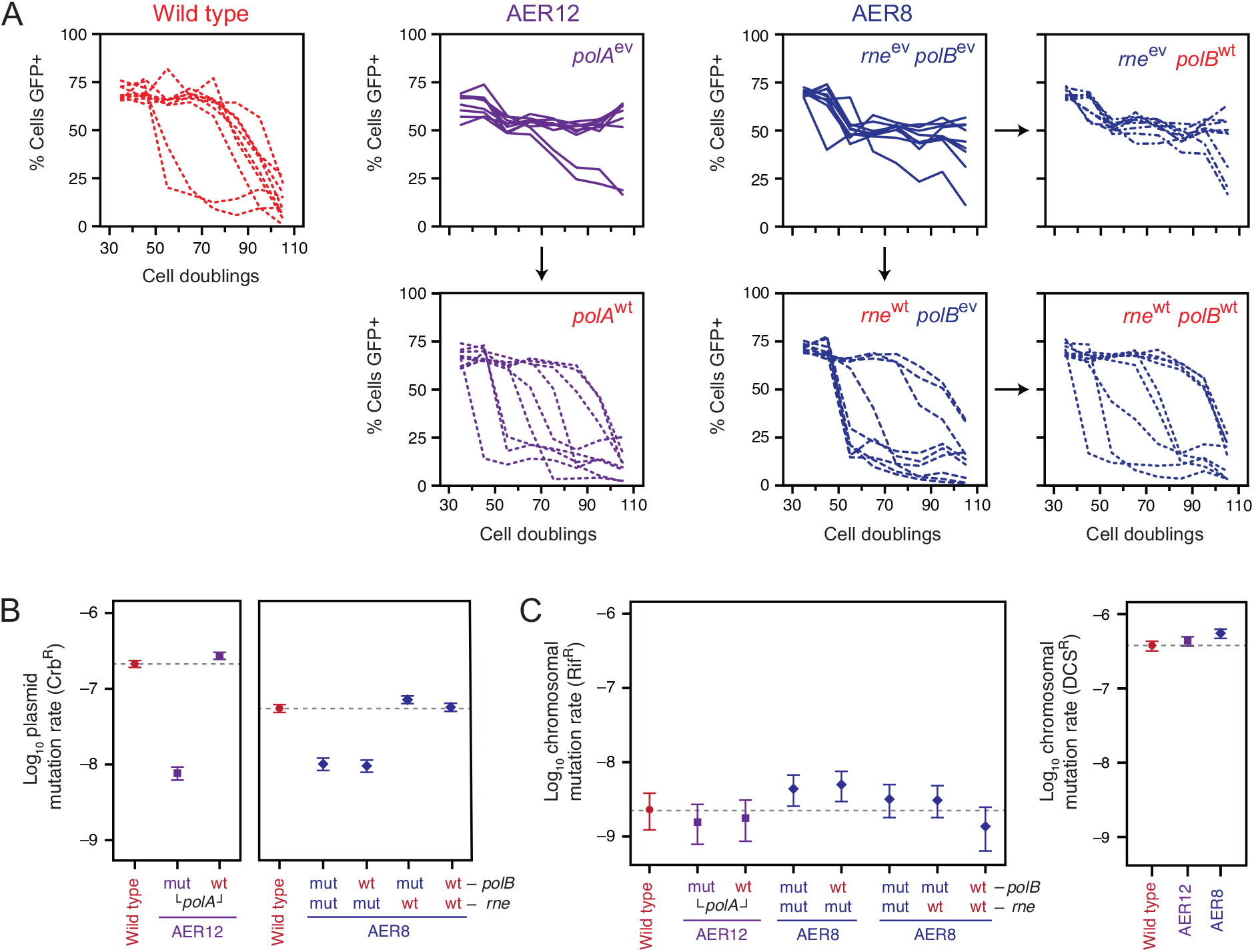
Evolutionary stability and mutation rates in PResERV and reconstructed strains. (**A**) Wild-type, evolved, and reconstructed strains were transformed with plasmid pSKO4. Nine independent populations were initiated from single colonies of each strain. Then, the prevalence of GFP-expressing cells within each population was monitored by flow cytometry over multiple daily serial transfers. Evolved strains are shown with solid lines. Dashed lines indicate that a strain contains one or more wild-type alleles, as indicated in red type. (**B**) Mutation rates to carbenicillin resistance (Crb^R^) due to reversion of a stop codon in the TEM-1.D254tag plasmid. Wild-type and evolved strains with mutant *polA, polB* and *rne* alleles (mut) reverted to wild-type alleles (wt), individually or in combination, were examined. Strains related to the evolved clone with a *polA* mutation (AER12) were tested in one experiment (left panel), and strains related to the evolved clone with *polB and rne* mutations (AER8) were tested in another experiment (right panel). (**C**) Mutation rates to rifampicin resistance (Rif^R^) (left panel) and D-cycloserine resistance (DCS^R^) (right panel) due to mutations in the *E. coli* chromosome. Each set of mutation rate measurements included the wild-type *E. coli* ancestor for comparison (dashed horizontal lines). Error bars are 95% confidence intervals estimated by the likelihood ratio methods from Luria-Delbrück fluctuation tests (**see Methods**).

We next used Luria-Delbrück fluctuation tests (26) to determine if the increase in evolutionary reliability in these strains was associated with a decrease in mutation rates. We first measured the rates of point mutations that reverted a stop codon in a β-lactamase gene cloned into another pBR322-based plasmid (20). Mutation rates to carbenicillin resistance, which requires mutating this stop codon to a sense codon, were significantly lower in each of the two focal evolved clones compared to wild type (**Fig. 4B**). In agreement with the GFP stability measurements, reversion of either the *polA* or *rne* mutation raised the mutation rate to that of the wild-type *E. coli* strain, and reversion of the *polB* mutation had no detectable effect on the mutation rate. We also measured chromosomal mutation rates in two further sets of fluctuation tests, selecting either for mutants resistant to rifampicin or to D-cycloserine. We did not find any significant improvements versus the wild-type strain in these assays (**Fig. 4C**). Thus, it appears that PResERV discovered *E. coli* mutants that primarily display lower plasmid mutation rates, with much less of an effect, if any, on mutation rates in the chromosome.

### Plasmid copy number and GFP fluorescence in evolved strains

Given previous reports on the effects of *rne* loss-of-function mutations (32), we were concerned that a reduction in ColE1 plasmid copy number in the PResERV evolved cells could give a false signal of improvement in our two assays. First, it would lower the GFP expression burden on host cells and thereby increase the number of cell doublings it would take for mutants to outcompete cells with the wild-type plasmid (i.e., it would increase the apparent evolutionary stability). Second, with fewer plasmids per cell there would be a smaller chance that any given cell would experience a mutation in one of its plasmids would lead to resistance in the stop codon reversion assay (i.e., it would reduce the apparent mutation rate per cell). Therefore, we measured the copy number of the pBR322 plasmid used in the mutation rate assay in the two focal evolved clones and four reconstructed strains using qPCR (**Fig. 5A**), and we also examined the per-cell GFP fluorescence from the pSKO4 plasmid in each strain (**Fig. 5B**).

**Figure 5.**
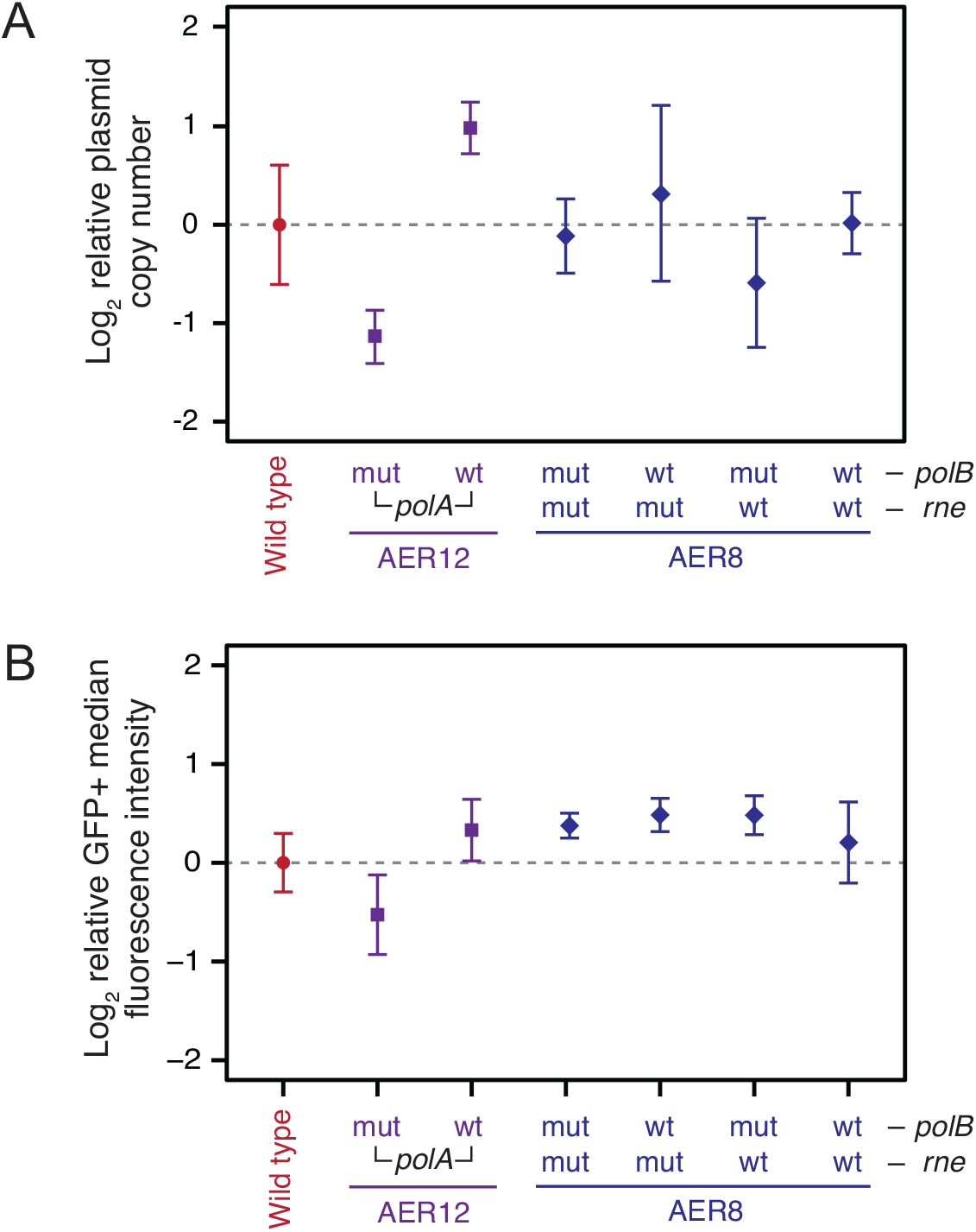
Plasmid copy number and GFP fluorescence in evolved and reconstructed strains. (**A**) Plasmid copy number for wild-type, evolved, and reconstructed strains determined by qPCR. Wild-type and evolved strains with mutant *polA, polB* and *rne* alleles (mut) reverted to wild-type alleles (wt), individually or in combination, were tested. The horizontal dashed line indicates the estimated copy number in the wild-type strain. Error bars show the standard error of the mean on log-transformed values from three biological replicates. (**B**) Initial GFP fluorescence of wild-type, evolved, and reconstructed strains as measured by flow cytometry. The median per-cell fluorescence intensity of the GFP+ subpopulation of cells was determined for each of nine replicate cultures immediately after outgrowth in liquid culture (after ~35 cell doublings). The graphed values are the log-averaged values of these medians. Error bars are 95% confidence intervals calculated assuming the logarithms of the medians are normally distributed. The horizontal dashed line shows the value for the wild-type strain.

The *polA* mutation did seem to reduce plasmid copy number somewhat. The evolved *polA* strain (AER12) had marginally fewer plasmids per chromosomal DNA copy when compared to wild-type (*p* = 0.0666, one-tailed *t*-test) and also exhibited reduced GFP fluorescent intensity (*p* = 0.0142, one-tailed *t*-test on log-transformed GFP+ subpopulation medians). Interestingly, other mutations in the evolved strain appeared to counteract the effects of the *polA* mutation, as reverting just this mutation to the wild-type allele increased both copy number (*p* = 0.0011) and GFP intensity (*p* = 0.0003). GFP signal (*p* = 0.0108) and perhaps copy number (*p* = 0.0893) were even greater in this *polA* revertant that still contained all other evolved mutations than they were in the original wild-type *E. coli* strain. Overall, these results suggest that there is a trade-off in the evolved *polA* mutant between plasmid copy number and mutation rate, but the reduction in plasmid copy number aof ~2-to 4-fold is much less than the ~30-fold reduction in mutation rates.

In contrast, plasmid copy number was unchanged from wild-type in the evolved *rne polB* strain (AER8) and all three reconstructed strains reverting those mutations singly and in combination (one-way ANOVA, *F*_4,10_ = 0.4393, *p* = 0.7777). Here, too, there was evidence that other mutations in these strains may have increased GFP intensity, as all four of the AER8-derived strains considered together had indistinguishable fluorescence intensities (one-way ANOVA, *F*_4,10_ = 0.4393, *p* = 0.7777) that were, as a group, significantly greater than that of the wild-type (*p* = 0.0112, one-tailed *t*-test). Therefore, the *polB* and *rne* mutations reduced plasmid mutation rates by ~6-fold with no detectable trade-off in terms of plasmid copy number or gene expression.

### Conclusions and Outlook

Natural selection has optimized the mutation rates of organisms to balance the deleterious effects of most mutations with the need for at least some supply of new mutations to fuel adaptive evolution (34). Evolution is unable to further reduce mutation rates in natural populations once the strength of selection against the deleterious genetic load from mutations is too weak to overcome genetic drift (35). We have shown that by artificially imposing strong selection for cells that do not give rise to mutations in a burdensome plasmid that we can overcome this ‘drift barrier’ and isolate *E. coli* that have lower-than-natural plasmid mutation rates. The antimutator alleles in these strains may be useful for constructing improved *E. coli* hosts for cloning and protein overexpression applications.

Despite years of studying the mechanisms of DNA repair and replication, we do not know the fundamental physiological limits on mutation rates that could potentially be achieved by tuning or augmenting these processes. Some studies have identified antimutator polymerase variants (33) or housecleaning proteins that can be overexpressed to reduce mutation rates (36). Often, however, these genetic studies are conducted in cells that are defective in DNA repair, so the mutations that they identify may only suppress a deficit, rather than improving upon already low mutation rates in a wild-type cell. Future applications of PResERV with the burdensome reporter gene encoded in the chromosome, rather than on a plasmid, might enrich for host cell variants that have a higher fidelity for replicating all components of the bacterial genome. This approach could also potentially be generalized to other cell types used for industrial bioproduction, such as yeast or Chinese hamster ovary cells. Ultimately, lowering mutation rates to arrest evolution can improve the foundations of synthetic biology so that engineered cells will function more predictably and reliably.

## DATA AVAILABILITY

Genome sequencing files have been deposited with the NCBI Sequence Read Archive under accession number SRP090775.

## ACKNOWLEDGEMENT

We thank Drew Tack and Andy Ellington for providing plasmid pTEM-1.D254tag and acknowledge the Texas Advanced Computing Center (TACC) at The University of Texas at Austin for providing high-performance computing resources.

